# Shared Molecular Signatures and Pathways in Post-Traumatic Stress Disorder, Schizophrenia, and Bipolar Disorder: An Integrative Transcriptomic and Machine-Learning Study

**DOI:** 10.64898/2026.02.26.708243

**Authors:** Mahfuj Khan, Md. Aminul Islam, Md. Arju Hossain

## Abstract

**Objective:** Psychiatric disorders such as post-traumatic stress disorder (PTSD), schizophrenia, and bipolar disorder (BD) are complex conditions with overlapping molecular features that remain poorly understood. The purpose of this study was to determine shared molecular abnormalities and potential therapeutic targets associated with the onset of these disorders.

**Methods:** RStudio (version 4.4.1) was used to process and analyze gene expression datasets from NCBI database in order to find differentially expressed genes (DEGs). FunRich, DAVID, STRING and NetworkAnalyst were used to gene ontology and pathway networks, protein– protein interaction, transcription factors and microRNAs and protein–chemical interactions. The discriminative ability of prioritized hub genes and transcription factors was evaluated by ROC analysis, and four machine-learning algorithms (Logistic Regression, Random Forest, SVM-RBF, and Gradient Boosting) were compared using leave-one-out cross-validation on the GSE403628 dataset (14 controls, 14 cases).

**Results:** Among 32 common genes, we identified several strong candidate hub proteins in significant pathways, including HLA-DRA, HLA-A, HLA-B, HLA-DOB, and BRD2. We also identified transcription factors (FOXC1, NFYA, RELA, GATA2, FOXL1, SRF, and NFIC), miRNAs (hsa-mir-129-2-3p, hsa-mir-148b-3p, hsa-mir-196a-5p, hsa-mir-26a-5p, hsa-mir-27a-3p, hsa-mir-23b-3p, hsa-mir-500a-3p, hsa-mir-423-5p, hsa-miR-142-5p, and hsa-miR-671-5p), and chemicals (Estradiol, Antirheumatic Agents, Valproic Acid and etc.). ROC analysis showed the highest AUC for HLA-DRA (0.760) among hub genes and FOXC1 (0.658) among transcription factors. In machine-learning comparisons, the hub-gene set consistently outperformed the transcription-factor set across all four algorithms, with Logistic Regression achieving the highest AUC (0.800). Under leave-one-out cross-validation, Random Forest and SVM-RBF yielded the highest accuracy (0.679).

**Conclusion:** Our computational approach identified potential biomarkers linking PTSD, schizophrenia, and BD to autoimmune inflammation and infectious disease pathways. ROC and machine-learning validation confirmed HLA-DRA as the strongest discriminator between disease and control samples, offering candidate targets for future investigation.

## 1. Introduction

Post-traumatic stress disorder (PTSD) is a diagnostic condition defined by involuntary and intrusive cognitive processes. People with PTSD often experience flashbacks, nightmares, and unwanted memories of the traumatic event ^1,2^. The condition also involves attention being drawn toward detecting dangerous triggers, along with difficulties in attention and memory ^3^. Because memory and attention problems are so common in trauma-affected individuals, it makes sense that researchers have turned to information-processing theories to explain PTSD ^3^. Over the past two decades, there has been a growing number of theoretical papers that try to explain worry through an information-processing lens. It is worth noting, however, that the DSM-5 no longer classifies PTSD as an anxiety disorder; instead, it now sits within its own category of trauma- and stressor-related disorders ^4,5,6^. As a result, anxiety-based theoretical frameworks may still have some bearing on PTSD research, but that connection is now more indirect and qualified than it once appeared ^6^.

Schizophrenia spectrum illnesses frequently exhibit concomitant symptoms that coincide with Schizophrenia spectrum illnesses frequently present with accompanying symptoms that overlap with several mental syndromes typically diagnosed in childhood, including inattention, restlessness, irritability, anxiety, and sadness. These symptoms closely resemble many of the side effects associated with schizophrenia and may therefore lead to a diagnosis of other psychiatric disorders during childhood and adolescence before schizophrenia is ultimately identified ^7^. Moreover, schizophrenia and certain pediatric psychiatric conditions may share common risk factors, including obstetric complications and risk genes, which may help explain some of the documented comorbidity ^8,9^. Furthermore, it has been proposed that autism, attention-deficit/hyperactivity disorder, and schizophrenia may share a neurological basis ^9,10^. Additionally, persistent neuroinflammation may serve as a shared risk factor for psychiatric disorders in children and adolescents as well as for late-onset schizophrenia ^11^. The initial presentation of schizophrenia may involve nonpsychotic syndromes or symptoms occurring during the prodromal phase or early stages of development, and may manifest as early as childhood ^8,12^.

Bipolar disorders refer to a group of persistent mental disorders, namely bipolar I disorder and bipolar II disorder. Bipolar I disorder is characterised by the occurrence of a syndromal manic episode, and pooled global estimates place the prevalence of bipolar spectrum disorders at approximately 0.74% ^13^. The diagnostic criteria for bipolar II disorder include the occurrence of a syndromal hypomanic episode and a major depressive episode. These estimates have mostly been obtained from research conducted in high-income nations. In developing and emerging economies, the reported lifetime rate of bipolar disorder has demonstrated variability; community surveys from Nigeria and Ethiopia, for example, indicate a lifetime prevalence of 0.1% to 1.83% ^14^. While certain persons with bipolar I illness may only have manic or mostly manic episodes, longitudinal research consistently indicates that depressive symptoms predominate over the long-term course, with patients spending roughly three times as many days depressed as manic or hypomanic ^15,16^. An often-reproduced observation in research on individuals with bipolar disorder is the early age at which symptoms present; systematic review evidence supports a trimodal age-at-onset distribution, with an early-onset subgroup averaging approximately 17 years and comprising roughly 45% of cases, and Global Burden of Disease 2021 analyses further indicate that incidence peaks at 15–19 years of age, with significantly higher rates in females than in males ^17^.

The specific connection between PTSD, Schizophrenia and Bipolar disorder remains unclear. Extensive study has been conducted on the relationship between these psychiatric disorders ^18,19,20^, with contradictory results. The contemporary absence of biomarkers to differentiate between these psychiatric diseases remains unresolved; no protein markers have been formally confirmed for either of these conditions. An essential aspect of analyzing transcriptomic data is the detection of biomarkers utilizing RNA-sequencing technology, namely by studying genes that exhibit differential expression ^21^. In recent studies, RNA-seq has been widely employed to ascertain illness pathophysiology, prognosis, and to discover novel biomarkers. The present work utilized RNA sequencing (RNA-seq) to examine transcriptional trends at the gene expression region ^21^. The aim of this study was to discover hub genes, microRNAs (miRNAs), and transcriptional factors (TFs) that could be used for personalized diagnosis and prognosis of post-traumatic stress disorder, Schizophrenia, and Bipolar disorder. Hence, the application of bioinformatics analysis to detect important genes, miRNAs, TFs, and related signaling pathways in relation to those disorders has immense potential for progression of future research in this domain.

### Graphical Abstract

The graphical abstract demonstrates datasets from PBMCs, neural progenitor cells, and human brain tissues was performed to identify differentially expressed genes, enriched pathways, hub genes, transcription factors (TFs) and microRNAs. Hub genes and TFs together with exploratory ROC and machine-learning classification across PTSD, schizophrenia, and bipolar disorder.

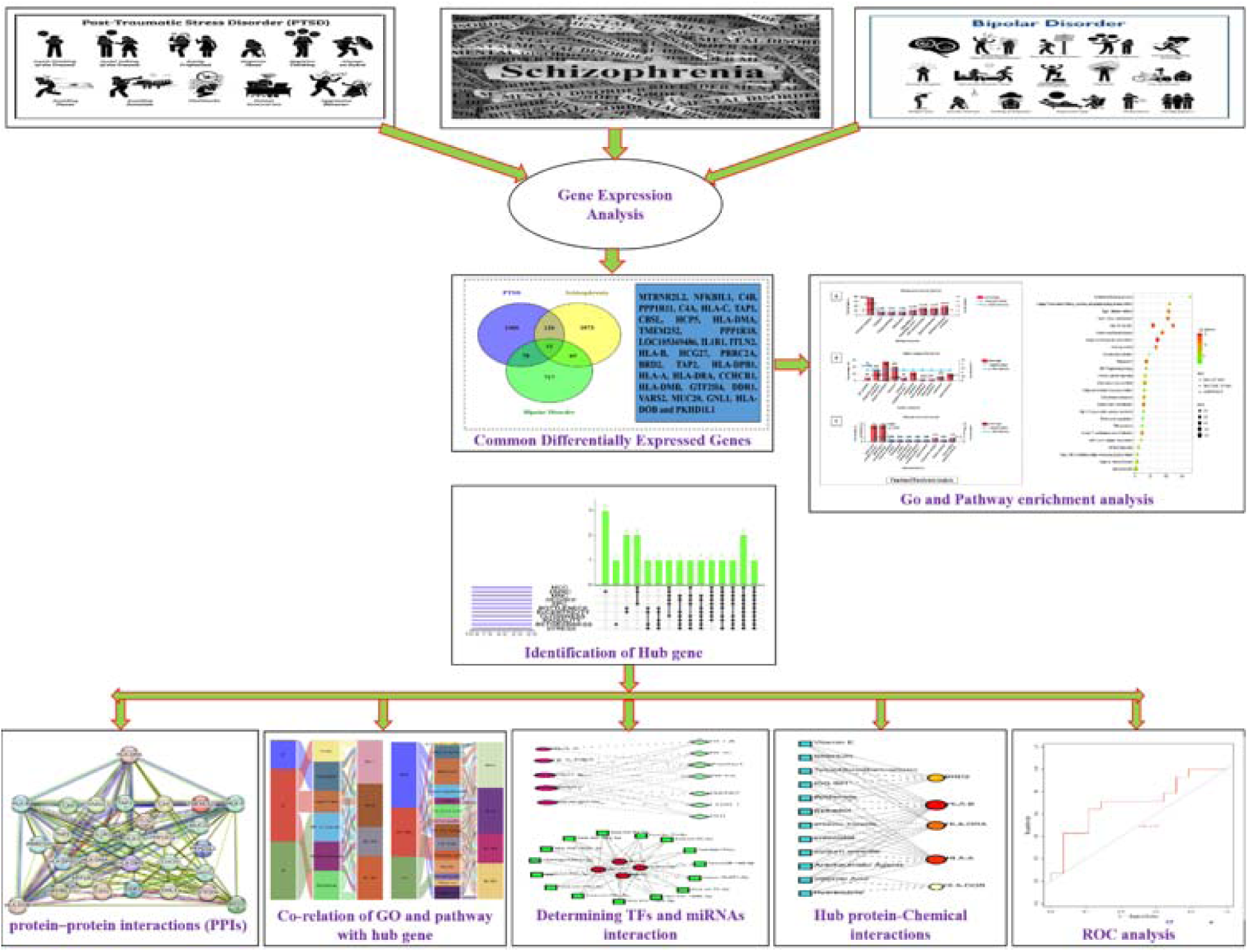

## 2. Materials and methods

### 2.1. Datasets used during the present research

(<u>NCBI</u>) GEO database ^22^ was employed in this investigation to evaluate the shared genomic interconnections across Post-Traumatic Stress Disorder, Schizophrenia, and Bipolar Disorder. The RNA-seq dataset employed in this study encompassed PTSD, Schizophrenia, and BD. The dataset on PTSD consisted of 5 advanced Blood transcriptomes PTSD samples and 5 normal samples (GEO accession ID: GSE8360 ^23^). These samples were analyzed using the high-throughput sequencing platform Illumina HiSeq. Compiled from brain progenitor cells, the Schizophrenia dataset (GEO accession ID: GSE119290 ^24^) consisted of 22 samples from cas persons and 22 samples from Schizophrenia patients. The Bipolar disorder dataset (GEO accession ID: GSE53239 ^25^) consists of 11 case samples and 11 control samples. The common platform GEO2R tool was utilized ^26^ for analysis the PTSD, Schizophrenia and BD. Hierarchical heat map of gene expression datasets of Post-traumatic stress disorder Schizophrenia and Bipolar disorder provided in Supplementary **Fig. S1 (A), S1(B) and S1(C)**.

### 2.2. Identification of DEGs and shared DEGs between PTSD, Schizophrenia and BD

The objective of differential expression analysis is to investigate the expression levels of genes under diverse situations. These genes offer biological understanding of the processes influenced by the specific condition of interest ^27^. The datasets were analyzed using R (version 4.4.1) to identify differentially expressed genes connected with PTSD, Schizophrenia, and Bipolar disorder, based on their corresponding control groups. Initially, we applied the log2 transform and statistical methods to standardize the gene expression data. To manage the rate of false discovery, we utilized the “Limma” package in the R language, using Benjamini-Hochberg approach ^28^. P-values below 0.05 and a |logFC| greater than 1 were used to identify the significant DEGs. The shared DEGs of GSE83601, GSE119290, and GSE53239 were identified using the Jvenn online VENN modeling program ^29^.

### 2.3. Functional enrichment analysis

We applied DAVID ^30^ and Funrich ^31^ employing Fisher’s exact test beside the combined DEGs to do functional enrichment assessment. By employing multiple datasets, we predicted the gene ontologies and signaling pathways associated with the predominant DEGs. The excessive appearance analysis revealed a set of signaling pathways and functional Gene Ontology terms that highlight the biological significance of the detected DEGs. Molecular pathways were proposed based on data from the databases KEGG ^32^ and Reactome ^33^ pathways, while gene ontologies were compiled for biological processes, cellular components, and molecular functions. Subsequent to the removal of redundant paths, only the pathways exhibiting statistical significance with a P-value below 0.05 are examined. Consequently, pathway-based analysis is an innovative approach to understand the relationships among complicated diseases through their underlying biological mechanisms. In contrast, the Gene Ontology is an organized framework utilized to delineate gene activities and their relationships with epigenetic modifications ^34^. The bubble plots were created using SRplot software ^35^.

### 2.4. Network assessment of protein–protein interactions (PPIs)

PPI network produced by shared differentially expressed genes was built using the <u>STRING</u> database (version 11.0) ^36^ to illustrate the structural and functional interactions among our designated DEGs and proteins. Protein-protein interactions in STRING illustrates the biological and functional connections between our specified DEGs and proteins. Using Cytoscape ^37^, PPI networks were seen among the DEGs. Eleven topological approaches, including MCC, MNC, Betweenness, Bottleneck, EcCentricty, DMNC, Degree, Closeness, EPC, Stress, and Radiality, were applied through the cytoHubba module available in Cytoscape to identify hub genes ^38^.

### 2.5. Correlation of most significant pathways and HubGs

An alluvial plot ^39^ illustrates the correlation between the name of process, the most significant pathways and potential hub genes. <u>SRplot</u> ^35^ is used to visualize the figure of alluvial plot. A complete selection of gene ontology feature, including the most significant pathways from the CC, BP, and MF, was shown in the GO alluvial plot. Furthermore, this diagram illustrates the 14 unique pathways obtained from the KEGG and Reactome pathways databases. The most enriched pathways together with their associated hub genes and log fold change (logFC) values are then entered into the SRplot web-based server, resulting in the creation of an alluvial graph illustrating the interconnections between the Gene Ontology keywords and the hub gene.

### 2.6. Determination of transcription factors and miRNAs

We applied NetworkAnalyst ^40^ tool for determining topologically feasible TFs binding with the hub genes in the JASPAR database ^41^. The relationships between miRNAs and their corresponding genes were examined to identify miRNAs that target gene expression to suppress protein synthesis ^42^. The principal databases for clinically confirmed miRNA-target relationships include Tarbase ^43^. Through the topological analysis of Network Analyst, we identified significant miRNAs from Tarbase.

### 2.7. Analysis of Protein-chemical compound

The screening of protein-chemical compounds facilitates the identification of individual chemical molecules responsible for protein interactions in comorbidities. Utilizing the advanced gene (hub gene) linked to PTSD, Schizophrenia, and Bipolar-disorder. Using the Comparative Toxicogenomics Database ^44^, We successfully identified protein-chemical interactions via Network Analyst ^40^.

### 2.8 Verification of the potential biomarkers

To evaluate how well the prioritized hub genes and transcription factors distinguished disease samples from controls, ROC-based and machine-learning analyses were carried out on the <u>GSE4036</u> dataset (28 samples: 14 controls and 14 cases). The normalized expression profiles of the selected molecular features served as input variables. Each hub gene and transcription factor was first assessed individually through receiver operating characteristic (ROC) analysis and the area under the curve (AUC).

#### 2.8.1 Receiver operating characteristic (ROC) analysis

The discriminatory performance of the selected molecular features was assessed individually by receiver operating characteristic (ROC) analysis ^45^. ROC curves and AUC values with 95% bootstrap confidence intervals (2,000 iterations) were derived from the normalized expression profiles of the selected hub genes and transcription factors through the SRplot online server (https://www.bioinformatics.com.cn/en). The corresponding area under the ROC curve (AUC) values were then calculated to evaluate their ability to identify disease versus control samples ^46^.

#### 2.8.2 Classification by Machine Learning

To compare the predictive performance of the two feature sets further, we evaluated four machine-learning algorithms: Logistic Regression (elastic net regularization, α = 0.5)^47^, Random Forest (100-200 trees, mtry = sqrt_features)^48^, Support Vector Machine with Radial Basis Function (SVM-RBF) kernel (C = 10, γ = 0.1)^49^, and Gradient Boosting (XGBoost, max_depth = 2-3, η = 0.1).^50^ To compare models, two feature sets were used such as Hub Genes and Transcription Factors.

#### 2.8.3 Cross-Validation with Leave-One-Out

To evaluate classification performance with the sample size at hand, leave-one-out cross-validation (LOOCV) was performed ^51^. For each iteration, one sample was omitted as the test sample, and the remaining samples were used to train models. This process was repeated until each sample had been the test sample. The predictions made were used to generate confusion matrices and calculate classification accuracy. The classification thresholds were individually optimized for each model (range: 0.3-0.7) to maximize accuracy^52^.

## 3. Results

### 3.1. Identification of DEGs and shared DEGs among psychiatric disorders

The Benjamini–Hochberg method was applied to each dataset to distinguish genes with lower expression from those with higher expression, thereby identifying viable candidates. The DEGs were identified in RNA-seq datasets using the particular conditions: absolute logFC values greater than 1 and less than -1 (logFC values >1 and <-1), and a significance level of P < 0.05. Specifically, genes with logFC >1 were regarded upregulated, whereas genes with logFC < -1 were regarded downregulated. The GSE83601 dataset revealed 1831 DEGs, the GSE119290 dataset revealed 1425 DEGs and GSE53239 dataset revealed 939 DEGs (**Table 1**). We employed the Venn diagram tool to identify shared differentially expressed genes among three datasets. A total of 32 prevalent DEGs were identified in (**Fig. 1**). On the other hand, **Table 1** indicates the datasets and quantitative outcomes of this investigation. We have categorized the study into PBMC, neural progenitor cells, and human brain to examine the variations in differentially expressed genes, as illustrated in the heatmap.

**Fig. 1:**
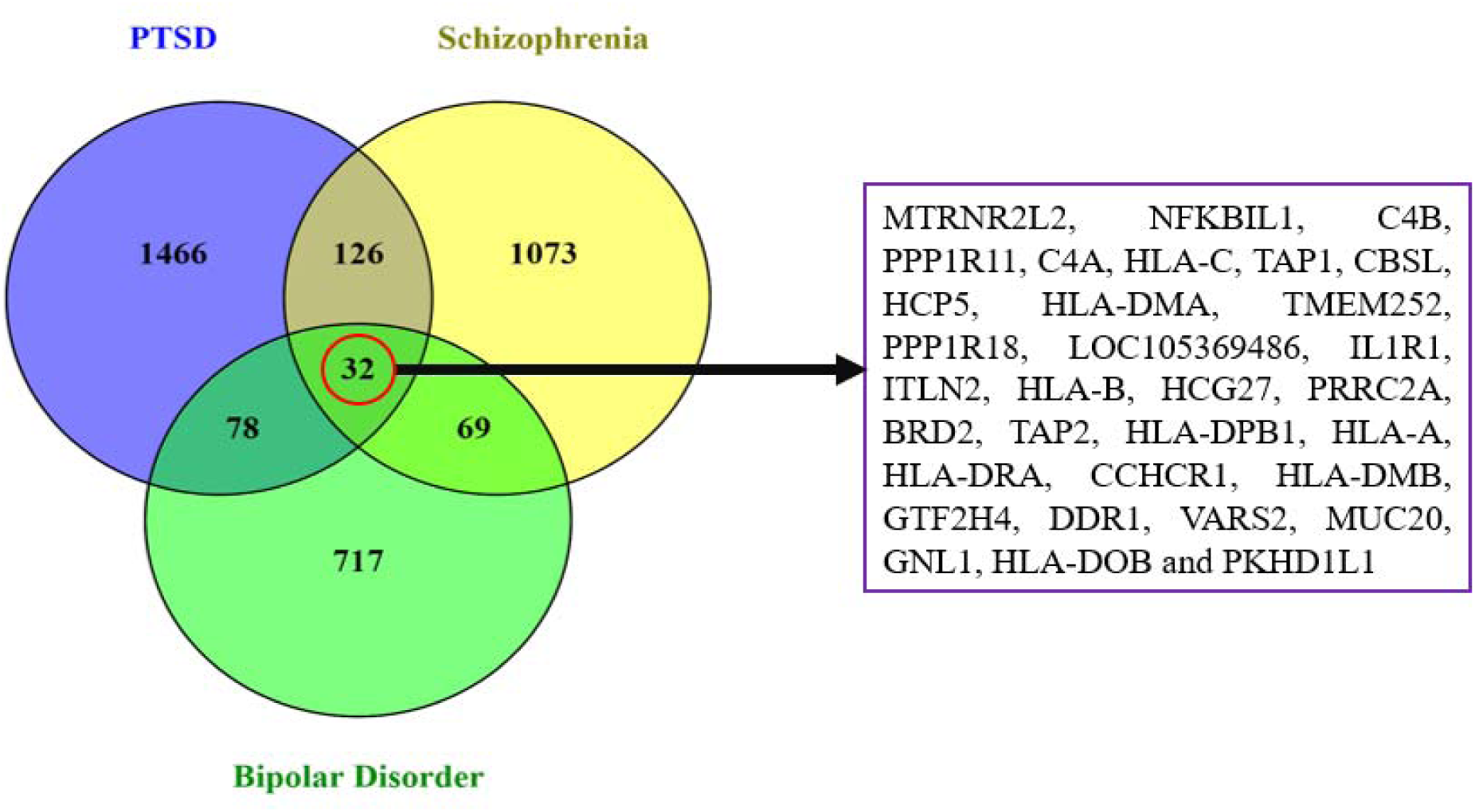
Shows the common DEGs between PTSD, Schizophrenia and Bipolar disorder. Red colors gene indicates the most significant in all disorders.

**Table 1:** Characteristics of GEO datasets and differentially expressed genes associated with three major psychiatric disorders: PTSD, schizophrenia, and bipolar disorder.

| **Disease name** | **GEO dataset** | **Cell type / tissue source** | **Case** | **Control** | **Total sample** | **Sequencing platform** | **Technology** | **Up DEGs** | **Down DEGs** | **Total DEGs** |
|---|---|---|---|---|---|---|---|---|---|---|
| PSTD | GSE83601 | PBMC | 5 | 5 | 10 | Illumina HiSeq 2000 | RNA-seq | 441 | 1390 | 1831 |
| Schizophrenia | GSE119290 | neural progenitor cells | 22 | 22 | 44 | Illumina HiSeq 2500 | RNA-seq | 748 | 677 | 1425 |
| Bipolar disorder | GSE53239 | human brain | 11 | 11 | 22 | Illumina HiSeq 2000 | RNA-seq | 839 | 100 | 939 |

### 3.2 Functional enrichment analysis

This work used Funrich to conduct gene ontology and DAVID database used for pathway enrichment analysis in order to uncover the biological importance and significant pathway associated with the shared DEGs studied. Gene ontology refers to the comprehensive collection of computable knowledge resources that consider gene functions and their components. An ontology formalizes a collection of knowledge, conceptually inside a specific framework. Both ontology and annotation serve the purpose of constructing a comprehensive biological structural model that is generally used to support biological applications. An examination of the gene ontology was conducted in three categories: biological process, cellular component, and molecular function. The GO database was used as the source for annotation. A summary of the top category of every database is presented in **Fig. 2(A-C)**. The comprehensive ontological examination for each type is also linearly characterized in the bar graph.

**Fig. 2:**
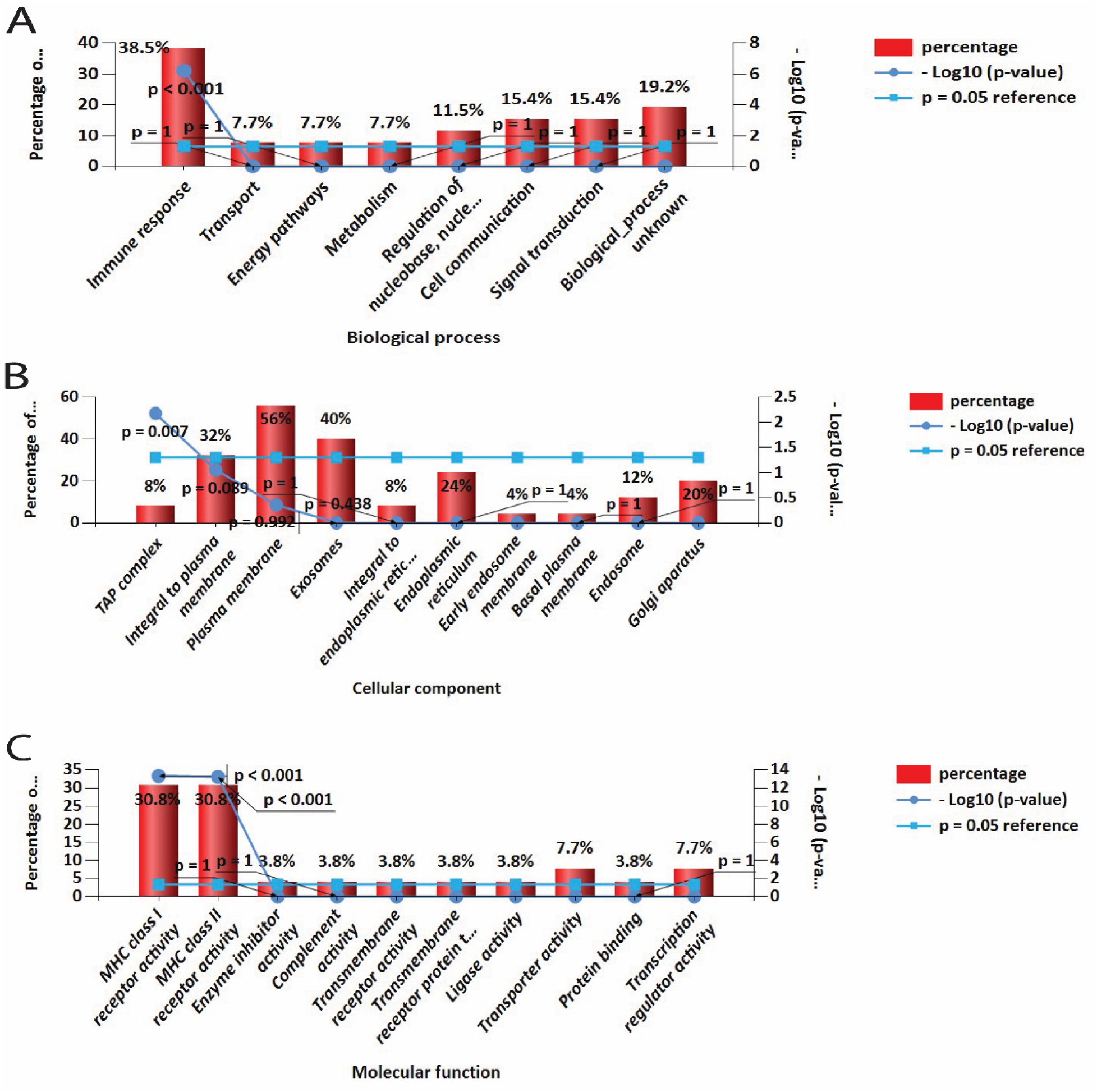
Bar diagram depicts the analysis of important GO pathways for diseases. (A) Biological process, (B) Cellular components and (C) Molecular functions. The top significant pathways selected based on a p-value < 0.05.

On the other hand, pathways analysis investigates how the organism responds to its intrinsic alterations. This approach is a paradigm used to illustrate the interplay of different diseases by means of fundamental molecular or biological mechanisms. From three global databases, namely KEGG, Reactome, and Wiki Pathways, we collected the most enriched pathways of the common DEGs in PTSD, schizophrenia, and bipolar disorder. The highest-ranking pathways derived from the chosen datasets. For further precision, the pathway enrichment analysis was also depicted in the multi group bubble plot provided in **Fig. 3**.

**Fig. 3:**
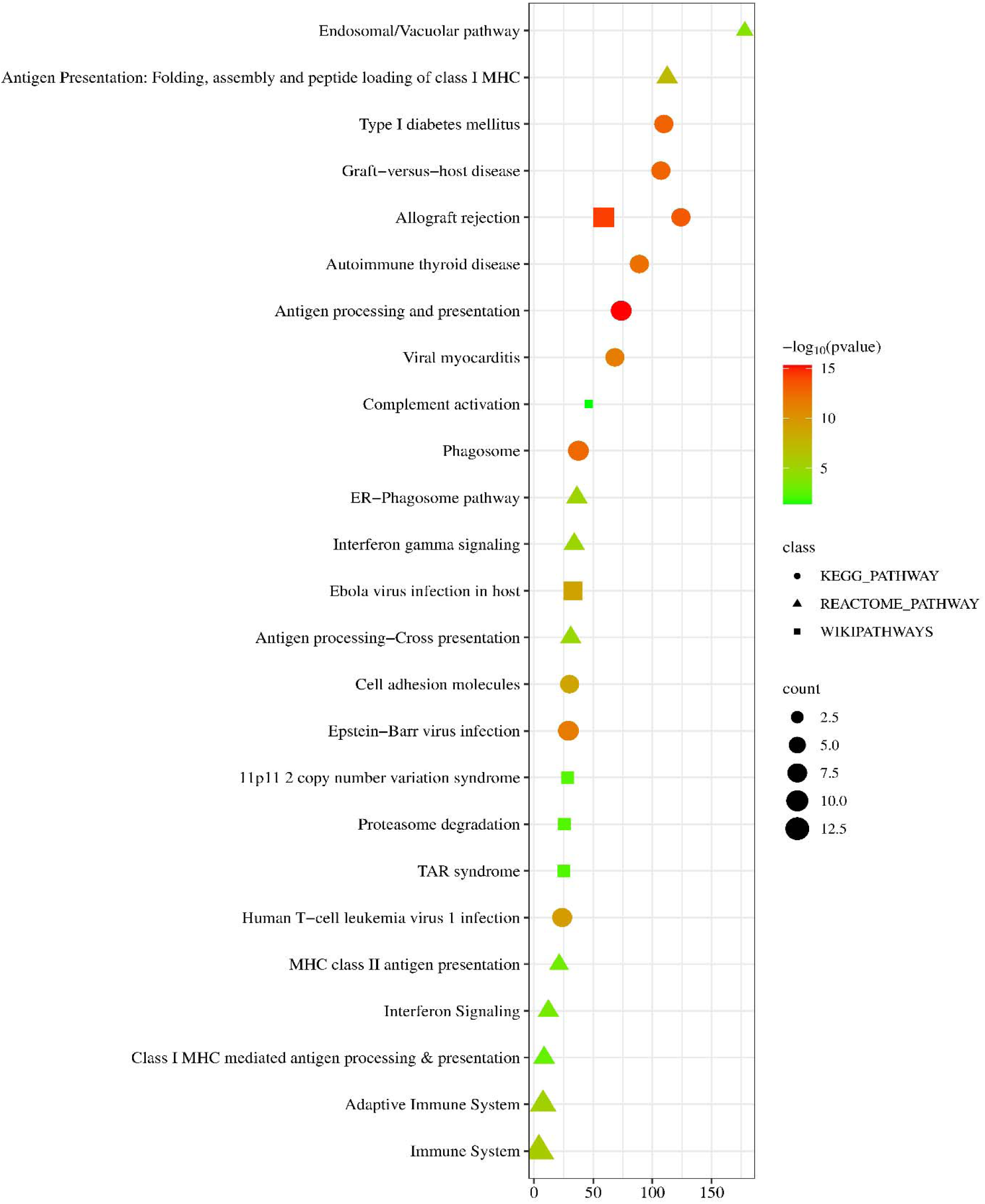
Hub gene enrichment results for the top metabolic pathways are visualized in a bubble plot using KEGG, Reactome, and Wiki Pathways data.

### 3.3 Determination of hub proteins

We analyzed the protein-protein interaction network obtained from STRING and displayed it in Cytoscape to predict the interactions and adhesion pathways of shared DEGs. **Fig. 4A** illustrates the PPI network of shared DEGs, which comprises 29 nodes and 135 edges. Simultaneously, the majority of linked nodes are recognized as important regulators in a PPI network. Using the Cytohubba plugin in Cytoscape, we identified the top 5 differentially expressed genes (15.62%) as the most influential genes in the PPI network study. The designated hub genes are HLA-DRA, HLA-B, HLA-A, BRD2, and HLA-DOB. The hub genes possess the capacity to function as biomarkers and may inform the creation of innovative therapeutic strategies for the studied disorders. To achieve a more thorough comprehension of the close association and proximity of hub genes, we have constructed a part of the network utilizing the Cytohubba plugin. Here shows six algorithms MCC, DMNC, MNC, DEGREE, BETWEENNESS and BOTTLENECK among the 11 algorithms in **Fig. 4(B)**.

**Fig. 4:**
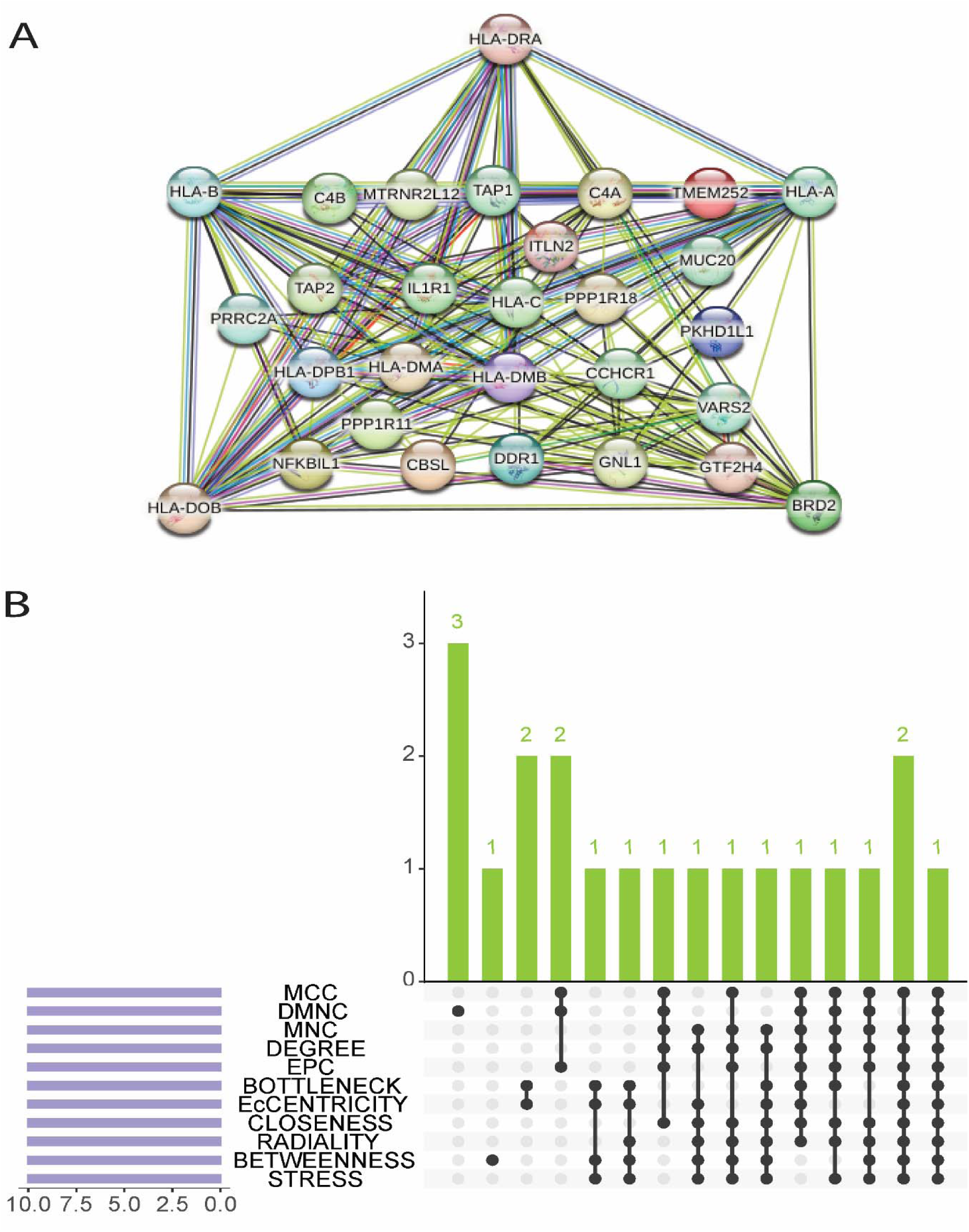
**(A)** Protein–protein interaction network of DEGs. Nodes positioned on the outer layer represent key genes (KGs), while the remaining circular nodes represent DEGs. (**B**) Hub genes were identified using the eleven topological metrics available in the cytoHubba plugin of Cytoscape, and the results are summarized in an UpSet plot. Notably, HLA-DRA was detected by all eleven algorithms.

### 3.4 Alluvial plot showing significant gene ontology and molecular pathway

The interconnections between the elements in an Alluvial plot, represented by radial geometric chords, serve to illustrate the interactions among the elements. Data groups are differentiated from each other by the use of distinct arc colors. The most enriched GO terms in each BP, CC, and MF category, including the top 6 most highlighted terms, are shown in **Fig. 5**. These terms are closely linked to the primary targets against PTSD, Schizophrenia, and Bipolar Disease. Conversely, the key 14 molecular pathways associated with the fundamental targets of PTSD, Schizophrenia, and Bipolar illness are illustrated in **Fig. 6**.

**Fig. 5:**
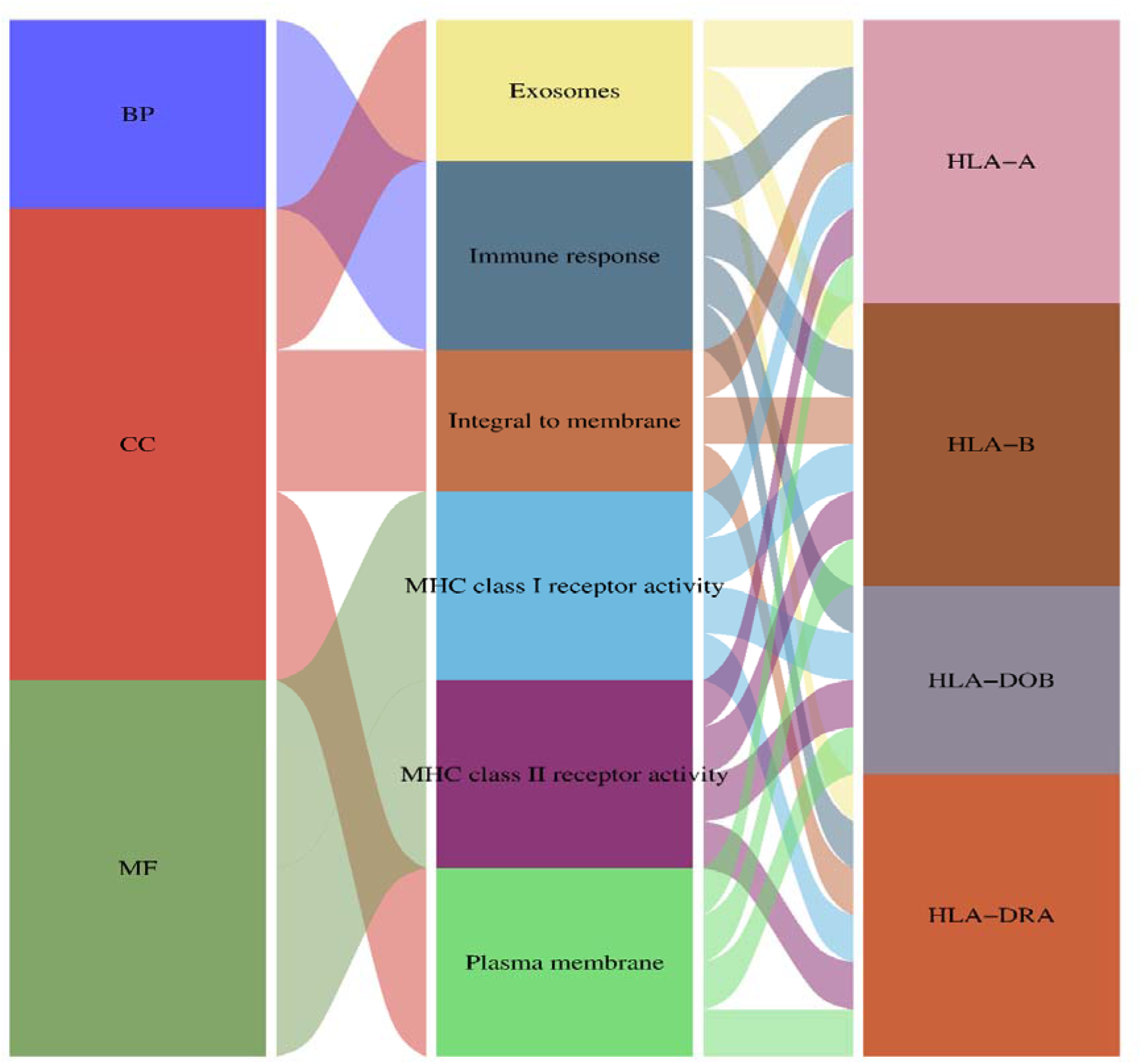
The alluvial plot of Gene ontology, go term and hub gene. The left column represents different go method, middle column represents GO term and the right represents hub gene, and the edge represents the relationship between them.

**Fig. 6:**
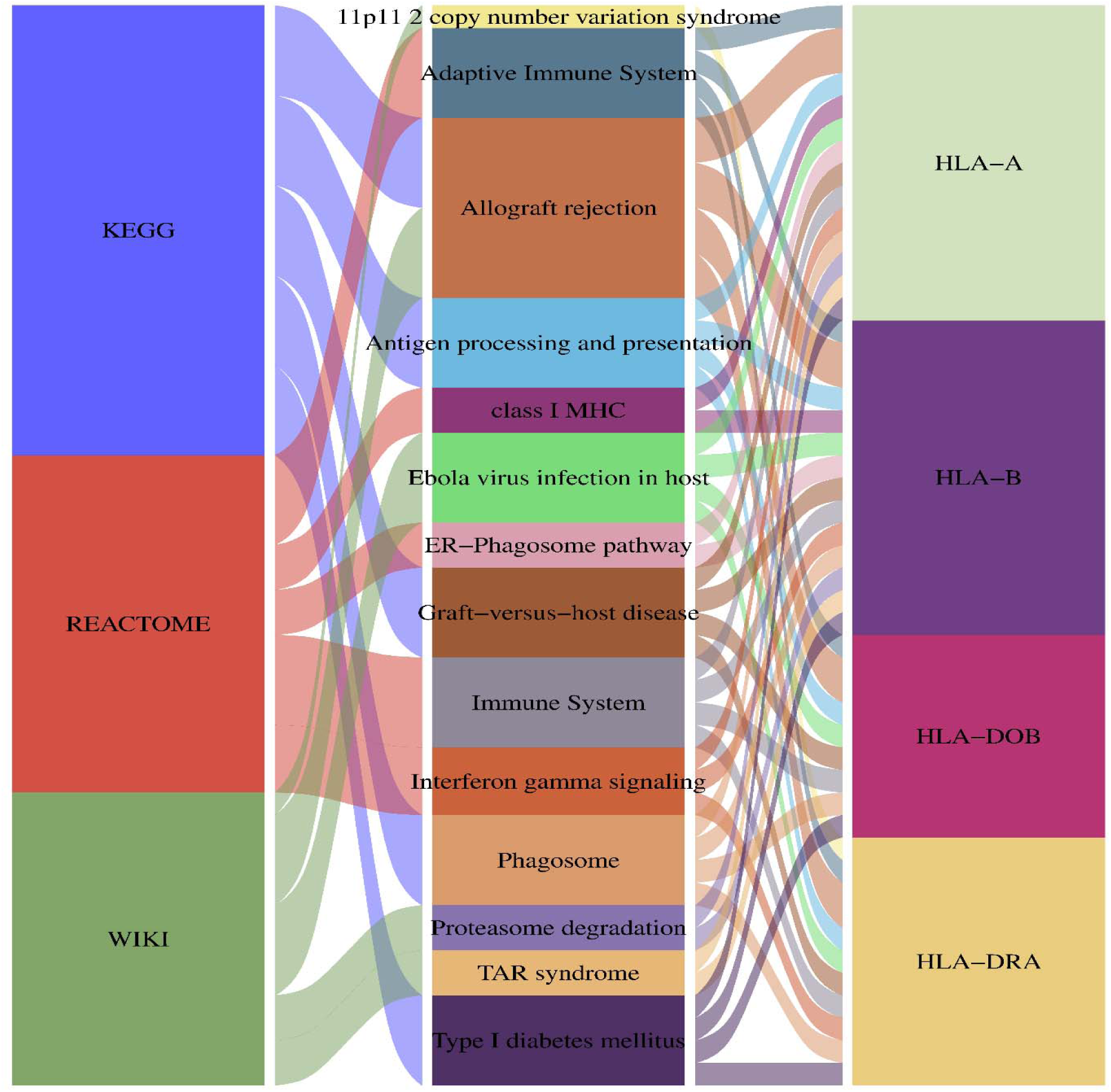
The alluvial plot of Pathways, pathway term and hub gene. The left column represents different pathway method, middle column represents pathway term and the right represents hub gene, and the edge represents the relationship between them.

### 3.5 Determination of regulatory signatures

In order to determine significant alterations occurring at the transcriptional level and gain understanding of the regulatory properties of the hub protein, we utilized a network-based method to decipher the regulatory transcription factors and microRNAs. The interaction of TF regulators with the hub gene is depicted in **Fig. 7A**. While **Fig. 7B** illustrates the correspondences among miRNA regulators and frequently occurring hub gene. Analysis of the gene interaction network involving transcription factors, hub genes, and microRNAs has determined that seven TFs and fifteen miRNAs regulate many common DEGs, indicating significant interaction among them.

**Fig. 7:**
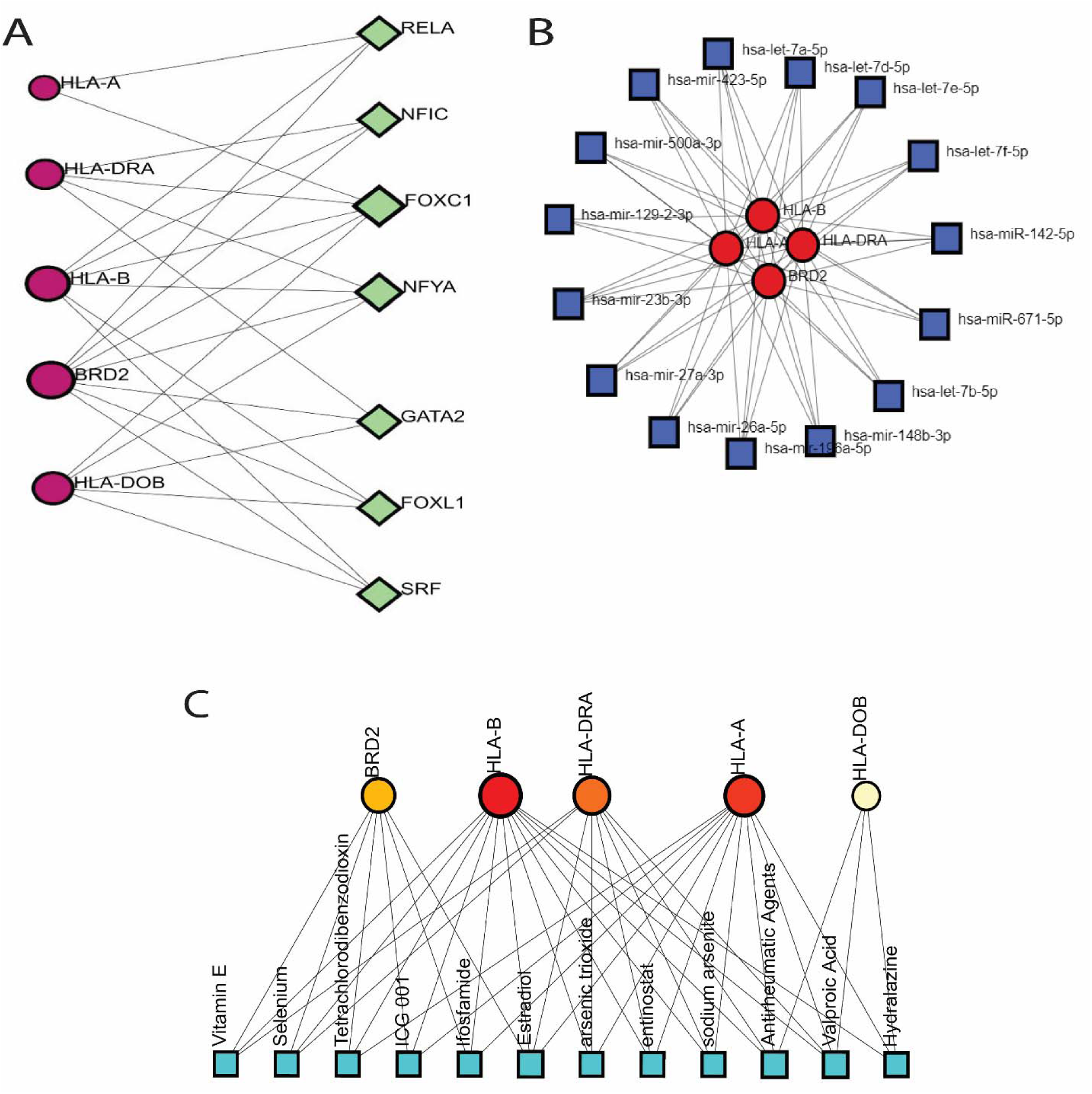
Gene regulator network analysis. **(A)** TFs and hub gene, **(B)** miRNAs and hub gene, **(C)** hub gene and chemical compound. Hub genes are shown as circle shapes, TFs are represented by diamond shapes, miRNAs and chemicals are represented by squared shapes.

### 3.6 The analysis of protein–chemical compounds interactions

Our studies have identified the protein-chemical interaction networks linked to PTSD, schizophrenia, and BD. A comprehensive assemblage of twelve chemical compounds has been identified, including Estradiol, Antirheumatic Agents, Valproic Acid, Selenium, Vitamin E, ICG 001, Ifosfamide, Tetrachlorodibenzodioxin, arsenic trioxide, entinostat, sodium arsenite, and Hydralazine. Their classification is that of highly enriched chemical agents. This network possesses the capacity to identify and classify significant proteins, such as HLA-DRA, HLA-A, HLA-B, HLA-DOB, and BRD2. The structures shown in **Fig. 7C** correlate to proteins and chemical agents, where circles symbolize proteins and squares identify chemical agents.

### 3.7 Validation of potential Biomarkers

#### 3.7.1 ROC (receiver operating characteristic) curve analysis for potential hub genes and transcription factors

Analysis showed that the selected molecular features differed in how well they discriminated between groups (**Fig. 8**). Within the hub-gene group, HLA-DRA had the highest AUC (0.760), followed by HLA-A (0.681), HLA-DOB (0.645), HLA-B (0.617), and BRD2 (0.551). Within the transcription-factor group, FOXC1 came out on top (AUC 0.658), followed by GATA2 (0.630), NFIC (0.615), FOXL1 (0.610), RELA (0.541), SRF (0.518), and NFYA (0.513). Taken together, the hub genes and transcription factors separated disease from control samples to varying degrees, with HLA-DRA and FOXC1 giving the best discrimination within their respective groups, while NFYA performed relatively poor. The hub gene (ROC) curve using the “pROC” package in R was used to examine the sensitivity and 1−specificity of recently discovered biomarkers hub gene are provided in Supplementary **Fig. S2**.

**Fig. 8.**
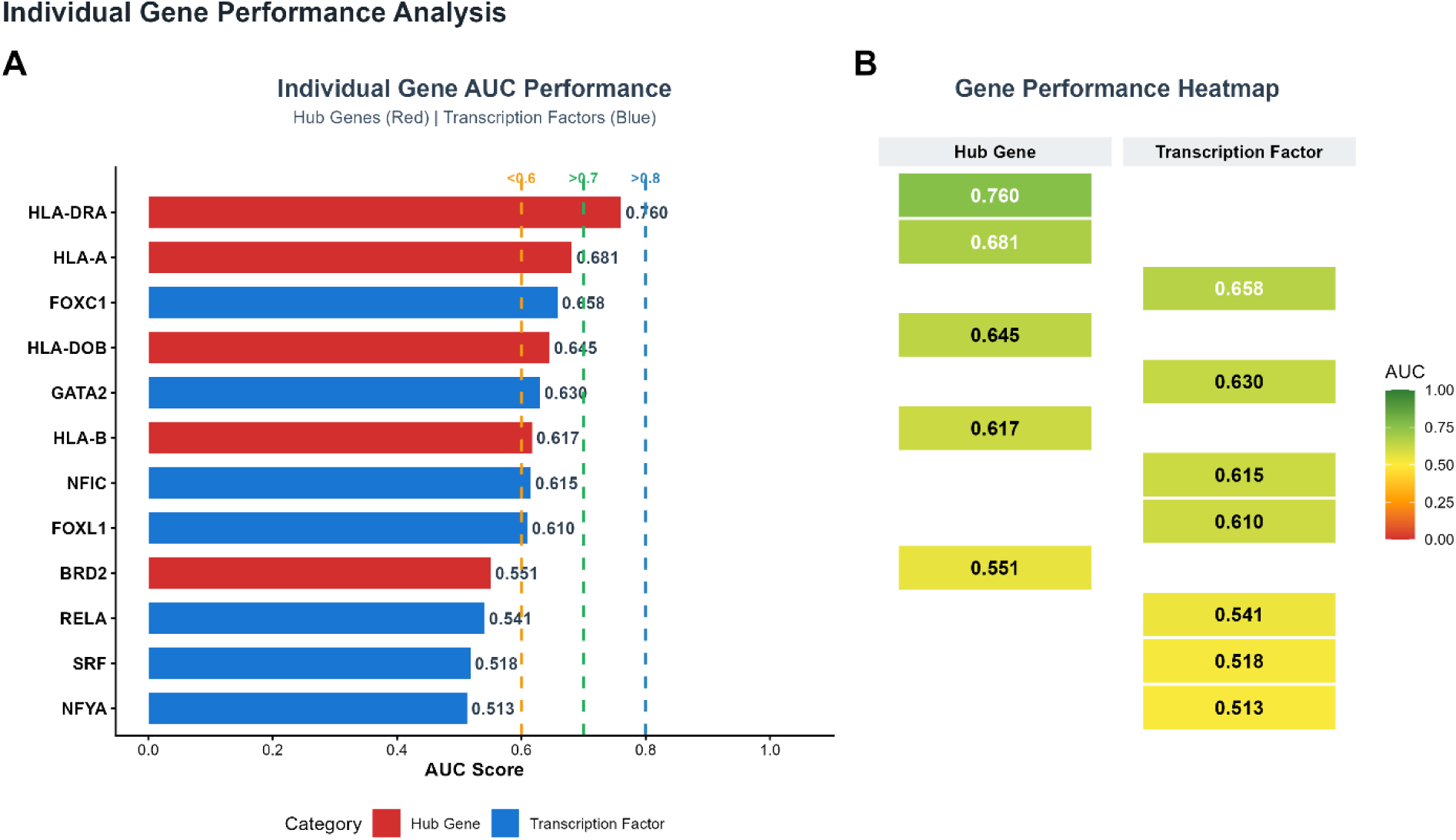
ROC analysis of the selected hub genes (HLA-DRA, HLA-B, HLA-A, BRD2, and HLA-DOB) and transcription factors (FOXC1, NFYA, RELA, GATA2, FOXL1, SRF, and NFIC). Random classification is indicated by the diagonal line.

#### 3.7.2 Machine-learning performance of hub genes and transcription factors

The machine-learning comparison revealed that classification performance differed between the hub-gene and transcription-factor (TF) feature sets across the four algorithms tested (**Fig. 9**). For the hub-gene set, AUC values were 0.800 with Logistic Regression, 0.762 with Random Forest, 0.758 with SVM-RBF, and 0.700 with Gradient Boosting; Logistic Regression was therefore the best-performing model for this set. For the TF set, the corresponding AUC values were 0.734, 0.738, 0.696, and 0.689, with Random Forest achieving the highest value (0.738). Overall, all four algorithms produced higher AUC values for the hub-gene set than for the TF set, and Logistic Regression using hub genes yielded the highest AUC overall.

**Fig. 9.**
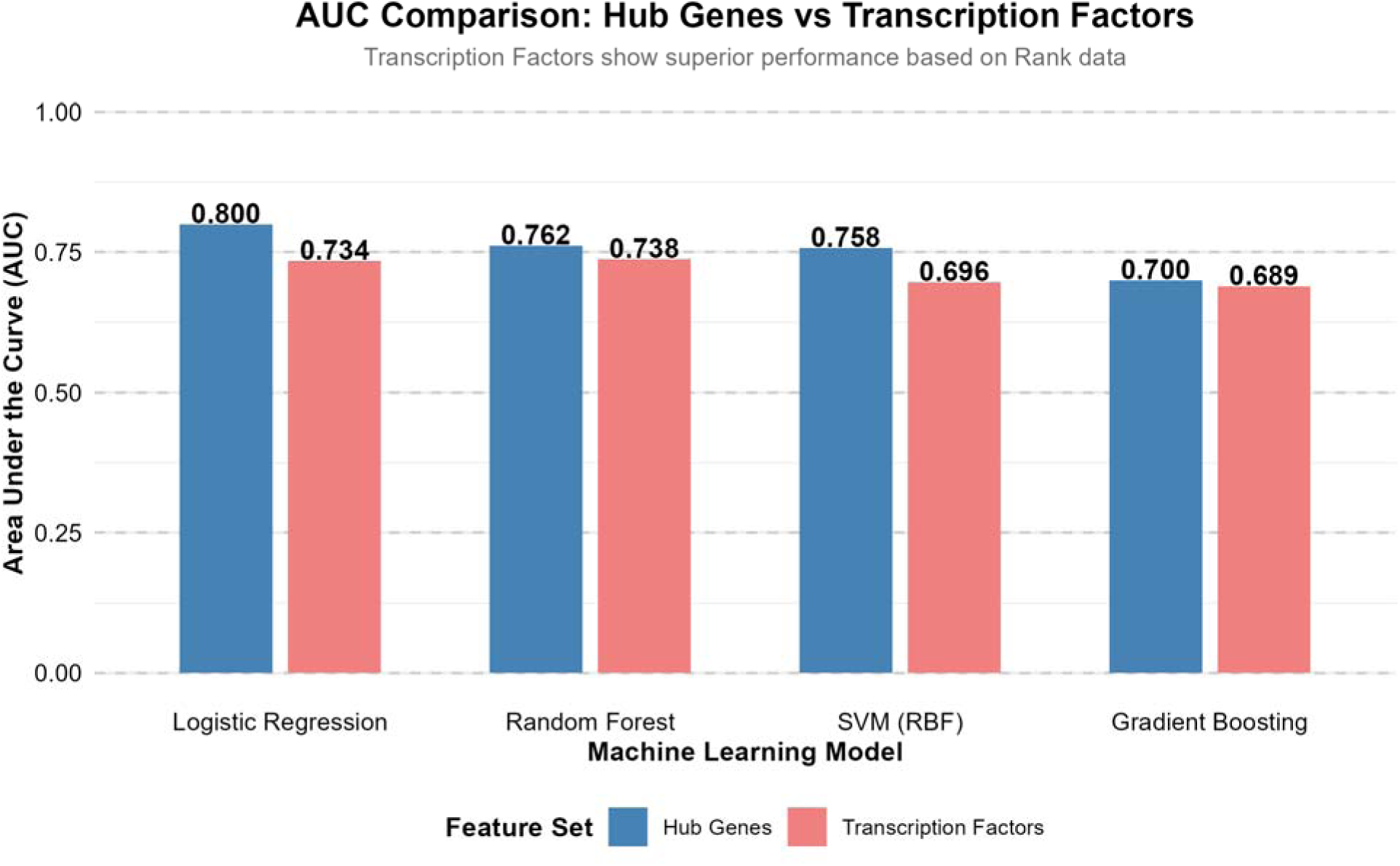
AUC-based comparison of the hub-gene and transcription-factor feature sets, illustrating the performance of Logistic Regression, Random Forest, SVM-RBF, and Gradient Boosting.

#### 3.7.3 Classification Performance under Leave-One-Out Cross-Validation

**Fig. 10** summarizes how the machine-learning models performed under leave-one-out cross-validation (LOOCV). Logistic Regression achieved an accuracy of 0.536, Random Forest 0.679, SVM-RBF 0.679, and Gradient Boosting 0.643. The top accuracy (0.679) was shared by Random Forest and SVM-RBF, with Gradient Boosting (0.643) and Logistic Regression (0.536) ranking below them. Taken together, Random Forest and SVM-RBF gave the strongest classification results, which agrees with their somewhat higher AUC values in the machine-learning comparison.

**Fig. 10.**
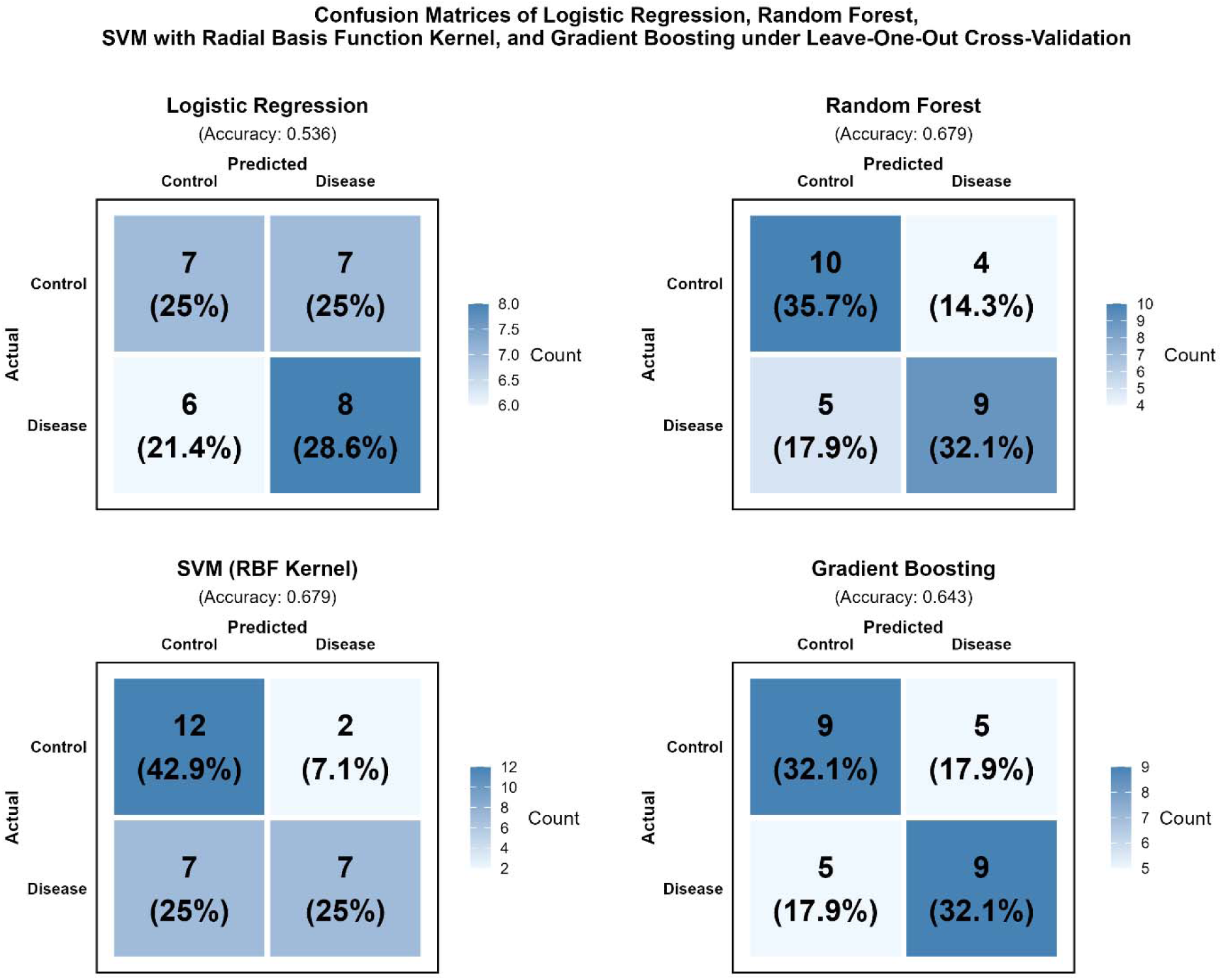
Classification results of Logistic Regression, Random Forest, SVM-RBF, and Gradient Boosting obtained under leave-one-out cross-validation, presented as confusion matrices.

## 4. Discussion

The present approach of diagnosing PTSD, Schizophrenia, and BD is based on the assessment of cognitive processes and the use of neuroimaging techniques. Nevertheless, there is a requirement for dependable and unique biomarkers to precisely detect and forecast the advancement of these psychiatric disorders. The current research utilized a systems biology methodology to comprehensively analyze gene expression patterns in several human cell types and brain regions linked to psychiatric disorders like PTSD, Schizophrenia, and BD. The present study has successfully discovered very promising molecular targets that have the ability to act as potential biomarkers for various psychiatric disorder. Therefore, our results may provide significant insights into the underlying mechanism of many psychiatric disorders.

Analysis of RNA sequencing datasets is intensively used in biological research and has become a valuable resource for discovering possible biomarkers ^53^. Gene expression profiling is often used to discover DEGs present in psychiatric disorders such as PTSD, Schizophrenia, and BD. This work sorted the gene expression dataset based on the p-value less than.05, |logF C| greater than 1.0, and |logF C| less than −1.0 to identify statistically significant DEGs that were up and down regulated.

The over-representation analysis identified Gene Ontology and molecular pathways elements related with the context of PTSD, Schizophrenia, BD, and neurodegeneration. The components comprising the GO category are plasma membrane, exosomes, immune response, MHC class I receptor activity, integral to membrane, and MHC class II receptor activity. Conversely, the analysis of metabolic pathways (KEGG, Reactome, and WiKi) indicates important genes that are associated with individuals with PTSD, Schizophrenia, and BD. The psychiatric disorders usually manifest the expression of specific genes related to allograft rejection, immune system, adaptive immune system, Type I diabetes mellitus, Ebola virus infection in the host and graft-versus-host disease.

Analysis of PPI reveals hub proteins responsible for transmitting signaling signals to other proteins in the networks. Using eleven topological metrics (HLA-DRA), hub genes were identified that have the potential to serve as biomarkers in psychiatric disorders. Furthermore, we have identified several other genes (HLA-B, HLA-A, BRD2, and HLA-DOB) that exhibit a robust correlation with PTSD, Schizophrenia, and Bipolar disorder, as established by a minimum of 9 Cytoscape algorithms. Impaired functioning of these genes can lead to the development of a serious psychiatric condition triggered by intense emotional, behavioral, and physical health issues. The presence of a HLA-DRA mutation has been demonstrated to induce Bipolar diagnosis ^54^. It has also been demonstrated as a potential gene associated with the disordered communication in PTSD and Schizophrenia ^55,56^ identified the HLA-DRA gene serves as an indicator in their investigation, which is closely linked to our study. Genetic abnormalities in the HLA-A gene may result in progressive neurological diseases. Multiple research studies suggest that a reduced HLA-A peptide-binding region is associated with the efficacy of psychiatric treatment in schizophrenia patients, as well as the emergence of PTSD and BD ^57,55,58,56^. Genetic mutations in the HLA-B gene can lead to Down syndrome, and research investigations have shown that individuals with Down syndrome often experience PTSD, schizophrenia, BD, as well as behavioral and mental disorders. So indirectly, the HLA-B gene may regulate PTSD, Schizophrenia, and Bipolar Disorder ^55,58,59^. HLA-DOB has been proven as a biomarker gene for PTSD, Schizophrenia and Bipolar disorder ^60,61^. BRD2 has been connected to the progression of Schizophrenia and BD origin in the cross-tissue examine ^62,63^, so we proved BRD2 as a biomarker pertinent to our research.

The excessive representation analysis identified genetic pathways and GO items in the areas of cellular component, biological process and molecular function related with neurodegeneration and PTSD, BD, and Schizophrenia. GO elements of utmost importance are plasma membrane, integral to membrane, exosomes, immune response, MHC class I & II receptor activity. Plasma membranes are predominantly expressed in several psychiatric disorders, BD and schizophrenia ^64^. MHC class I & II receptor activity associated with bipolar disorder ^65^. Immune responses: We documented the neuro-immune responses in euthymic bipolar illness patients submitted to the Trier Social Stress Test, noting significant reactivity associated with mental problems ^66^.

Metabolic pathways analysis using KEGG, WiKi, and Reactome identifies important genes associated with PTSD, BD, and Schizophrenia in patients. Allergic rejection, Type I diabetes mellitus, and the Immune System route are predominantly manifested in these psychiatric disorders. The involvement of allograft rejection in the evolution of BD and other psychiatric disorders ^67^. Type I diabetes mellitus: dysregulation of Type I diabetes mellitus play a vital role in progression of psychiatric disorders ^68^. Immune System: Increasing data indicates that components of the immune system significantly affect brain function, impacting several aspects of mental and neurological illnesses ^69^.

Our study also included an examination of the connection between Hub genes and TFs, as well as hub genes and microRNAs. Identified transcriptional regulatory transcription factors (FOXC1, NFYA, RELA, GATA2, FOXL1, SRF, and NFIC) in several psychiatric disorders correspond to our previous network-based approach for identifying overlapping molecular fingerprints in psychiatric disorders using RNAseq data.

Transcription factors FOXC1 (Forkhead Box C1) were identified in the regulatory network as potentially associated with BD and central cellular functionality. Hence, the disruption of the biomolecule FOXC1 could be utilized as potential biomarkers and curative targets in the development of treatments for PTSD, Schizophrenia, and BD ^70,71,72^. RELA exhibited a learning deficiency in the spatial variant of the radial arm maze ^73^, revealing the crucial role of the RELA gene in memory function, maybe associated with the underlying mechanisms of memory impairment in schizophrenia and other psychiatric disorders ^70,74,75^. A reduction in neuroglobin expression was seen upon the knockdown of GATA2 (GATA-binding factor 2). The genes GATA2 have been associated with PTSD, bipolar disorder and schizophrenia ^70,71,76^. Serum response factor (SRF) has been identified as a potential transcription factor that has a role in coordinating the MBR module in females. Upregulation of transcription factors linked with MBR Spinal reactivity factor (SRF) is a significant contributor to the PTSD-like reaction to stress in females ^70,77,78^. Conversely, prior research has shown a significant correlation between transcription factors FOXL1 (Forkhead Box L1) and NFIC (Nuclear factor IC) with PTSD, Schizophrenia, and BD, which we have elucidated with pertinent bibliography ^79,70,80^. Our investigation revealed a significant association between NFYA (Nuclear transcription factor Y subunit alpha) with PTSD, Schizophrenia, and BD, contrary to previous studies which did not establish such link.

Therefore, the dysregulation of miRNAs could serve as biomarkers. First ten miRNAs identified in the miRNA-Hub gene network were hsa-mir-129-2-3p, hsa-mir-148b-3p, hsa-mir-196a-5p, hsa-mir-26a-5p, hsa-mir-27a-3p, hsa-mir-23b-3p, hsa-mir-500a-3p, hsa-mir-423-5p, hsa-miR-142-5p, and hsa-miR-671-5p. Genetic mutations in the hsa-mir-129-2-3p gene are linked to psychopathologies that often develop in adolescents, including depression and schizophrenia.

These mutations are also highly connected with PTSD and Bipolar disorder, which pose a significant social risk ^81,82,83^. The identification of hsa-mir-148b-3p as a potential biomarker associated with cortical development, neuronal differentiation, neuron survival, and synapse formation indicates its alteration in PTSD, schizophrenia, and bipolar disorder ^84,82,83^. hsa-mir-196a-5p was scientifically proven PTSD, Schizophrenia and BD related miRNA ^85,82,86^. Differential expression of hsa-mir-26a-5p in PTSD, Schizophrenia, and BD has been established as a promising biomarker ^87,88,89^. The hsa-mir-27a-3p Although it may be tempting to hypothesise that targeting well validated miRNAs (mir-27a-3p) can have beneficial effects in people with BD, it is also closely linked to Schizophrenia and PTSD ^90,84,71^. Dysregulation of hsa-mir-23b-3p as Biomarkers in Psychiatric Disorders, was also observed in previous study where it connected with PTSD, Schizophrenia and BD ^91,92,71^. In spite of demonstrating minimal enrichment for bipolar disorder GWAS reported genes, the hsa-miR-142-5p-target gene set exhibited the topmost enrichment. Strong enrichment for brain imaging was not observed in any of the target gene sets. Genes identified by GWAS and associated with the development of PTSD and Schizophrenia should be considered as possible biomarkers in both disorders ^93,84^. miR-423–5p were differentially expressed in schizophrenia, it also a strong connection with PTSD and Bipolar disorder ^94,84^. Contrary to previous published findings, our investigation revealed a significant association between hsa-mir-500a-3p and hsa-miR-671-5p and PTSD, Schizophrenia, and BD. These chemicals affect genes at both the transcriptional and post-transcriptional stages.

Taking into account the importance of these key genes and their capacity to significantly influence the pathological processes in neurodegeneration associated with specific psychiatric diseases. A research investigation was undertaken to evaluate the connection among proteins and chemicals, with the aim of identifying molecules that may potentially influence these interactions. The interaction network revealed a total of 11 chemical compounds, including Estradiol, Antirheumatic Agents, Valproic Acid, Selenium, Vitamin E, ICG 001, Ifosfamide, Tetrachlorodibenzodioxin, arsenic trioxide, entinostat, and sodium arsenite.

The aforementioned statement implies that our approach have the capability to reveal fundamental pathways implicated in the progression of certain psychiatric diseases. In addition, it may generate novel hypotheses regarding the mechanisms that underlie these diseases and identify new biomarkers. Statistical analysis of genetic data will be essential in enhancing predictive medicine and elucidating the basic mechanisms connecting PTSD, Schizophrenia, and BD. Moreover, these investigations may reveal novel therapeutic targets. Therefore, further investigation is required to accurately evaluate the biological significance of the potential target possibilities identified in this work.

## Conclusion

By analyzing transcriptomic data from brain tissue and psychiatric disorder patients, we aimed to identify commonly differentially expressed genes. These shared DEGs were then used to examine related pathways, protein–protein interactions, transcription factors, and microRNAs. Through this approach, we identified potential biomarker transcripts that are commonly disrupted in psychiatric illnesses and proposed prospective pharmaceuticals specifically targeting the biomarkers we identified. To validate our findings, we employed ROC analysis and machine-learning classification as predictive and discriminative measures to support the diagnosis and treatment of PTSD, schizophrenia, and BD. The bioinformatics methodology used in this study may offer novel insights into possible targets implicated in susceptibility to psychiatric disorders and facilitate the development of drug-discovery programs aimed at treating these conditions.

## Acknowledgements

The authors would like to express their sincere gratitude to **Md. Arju Hossain**, Department of Biochemistry and Biotechnology, Khwaja Yunus Ali University, Sirajganj 6751, Bangladesh, for his valuable support and constructive suggestions. We also gratefully acknowledge the support of the **Center for Advanced Bioinformatics and Artificial Intelligence Research (CABAIR)**, Department of Computer Science and Engineering, Islamic University, for providing an encouraging research environment and technical assistance that contributed to the successful completion of this study.

## Ethical approval and informed consent statements

This research requires no approval from any ethics committee as this research uses data that is publicly available.

## Data availability statement

We have used publicly available data. Data for RNA-seq analysis were collected from NCBI. The Gene Expression Omnibus (GEO) database from NCBI was used to access the GSE83601, GSE119290, GSE53239 and GSE4036 datasets.

## Authorship contribution statement

**Mahfuj Khan:** Conceptualization, Methodology, Software, Validation, Formal Analysis, Writing-Original Draft, Writing – Review & Editing. **Md. Aminul Islam:** Data Curation, Formal Analysis, Visualization, Writing – Original Draft & Writing – Review & Editing. **Md. Arju Hossain:** Conceptualization, Methodology, Software, Validation, Supervision, Writing – Review & Editing.

## AI Disclosure Statement

The authors declare that no artificial intelligence (AI) tools or technologies were used in the writing, analysis, or preparation of this manuscript. All work was conducted solely by the authors.

